# The ecology of potentially pathogenic *Vibrio* spp. in a seagrass meadow ecosystem

**DOI:** 10.1101/2024.06.15.599152

**Authors:** Rebecca Gebbe, Katharina Kesy, Dorothea Hallier, Anne Brauer, Stefan Bertilsson, Matthias Labrenz, Mia M. Bengtsson

**Affiliations:** Institute of Microbiology, University of Greifswald, 17489 Greifswald, Germany; Institute of Biochemistry and Biology (IBB), University of Potsdam, 14476 Potsdam, Germany; Fraunhofer Institute for Celltherapy and Immunology, Branch Bioanalytics and Bioprocesses (IZI-BB), 14476 Potsdam, Germany; Department of Aquatic Sciences and Assessment, Swedish University of Agricultural Sciences, 75007 Uppsala, Sweden; Leibnitz Institute for Baltic Sea Research Warnemünde (IOW), 18119 Rostock-Warnemünde, Germany; Institute of Marine Biotechnology, 17489 Greifswald, Germany

**Keywords:** Seagrass microbiome, marine biofilms, *Vibrio vulnificus*, OneHealth, Baltic Sea ecology

## Abstract

Seagrass meadow ecosystems offer several valuable ecosystem services in coastal regions around the world. Recent studies have suggested that one such important service is reduction of pathogenic bacteria and specifically *Vibrio* spp. in adjacent waters. The specific mechanisms of pathogen reduction remain unclear, although increased sedimentation has been suggested as one likely process for pathogens to be quenched from the water column. Whether *Vibrio* spp. persist in the sediment or in other compartments of the seagrass meadow is currently not known, but it has been shown that marine surface biofilms can function as reservoirs of pathogenic vibrios. This general feature may also apply to seagrass and sediment surfaces. In this study, we investigated the relative abundance and community ecology of *Vibrio* spp. bacteria in Baltic Sea seagrass meadows using both culturing and culture-independent methods. While we did not detect a significant reduction of *Vibrio* spp., the highest relative abundances of *Vibrio* spp. were observed in the water column above unvegetated sites as compared to seagrass meadows. We also detected high relative abundances of *Vibrio* spp. on seagrass roots, supporting previous observations that marine surfaces are selectively colonized by *Vibrio* spp., implying that these habitats are important for the persistence and possibly release of *Vibrio* spp. into the water column. Our results emphasize the need to understand the interactions of pathogenic bacteria with coastal habitats, including interactions with host organisms such as seagrasses that provide biofilm microenvironments, in order to understand how diseases associated with these organisms develop.

## 1. INTRODUCTION

The genus *Vibrio* is a diverse group of marine bacteria which contains a remarkable number of opportunistic pathogens of marine animals and humans (Baker-Austin et al. 2018). There is rising concern about the prevalence of *Vibrio* infections due to the impacts of climate change, such as rising seawater temperatures which stimulate growth of several *Vibrio* pathogens (Baker-Austin et al. 2010, Vezzulli et al. 2013, Le Roux et al. 2015). As environmental opportunistic pathogens, vibrios are adapted to natural marine or brackish habitats by tolerating a wide range of salinity and temperature conditions while not primarily adhering to a pathogenic lifestyle. This could lead to coastal waters serving as a reservoir of these pathogenic bacteria and it is therefore important to understand the ecology of both pathogenic *Vibrio* spp. and related non-pathogenic strains to predict and mitigate health risks in marine environments. From a OneHealth perspective, human health and animal welfare must be understood in the context of environmental state (Zinsstag et al. 2011). *Vibrio* pathogens are a good example of how changing environmental state manifested as increasing coastal surface temperatures and eutrophication enhancing algal blooms may trigger the growth of these global pathogens (Eiler et al. 2007), which may then in turn have strong impacts on both marine wildlife and humans that use marine environments for recreation or consume marine food products (Marques et al. 2022).

The Baltic Sea is the largest brackish water body in the world, and is characterized by salinity gradients spanning marine to freshwater conditions across large geographic scales while also displaying local and seasonal fluctuations depending on the region (Kniebusch et al. 2019). In the south-eastern Baltic sea, e.g. the coasts of eastern Germany and Poland, the coastline is characterized by shallow lagoon systems, where salinity tends to be lower than on the open coast. In addition, surface water temperatures can exceed 15-20 °C for longer periods in these systems, especially during summer heatwaves (Baker-Austin et al. 2013). These specific conditions are risk factors for outbreaks of pathogenic strains of *Vibrio vulnificus* (Schütt et al. 2023), a species documented to cause severe infections with a high case fatality rate in mainly immunocompromised persons in this region (Dalsgaard et al. 1996, Ruppert et al. 2004, Baker-Austin et al. 2017). In Germany alone, 33 such infections were reported from 2018-2019 (Brehm et al. 2021) and there is currently an increasing trend with more frequent and longer lasting summer heatwaves (Bier et al. 2015, Vezzulli et al. 2016, Baker-Austin et al. 2017) which may exacerbate the problem. However, other less known factors may also play a role in stimulating or mitigating *Vibrio vulnificus* outbreaks. For example, biotic interactions with other marine organisms are key factors that structure marine bacterial communities and this also applies to *Vibrio* spp. which are sensitive to predation by bacterivorous protists and animals (Lutz et al. 2013). Conversely, *V. vulnificus* may infect, associate, interact and benefit from marine animals and primary producers such as algae and seagrass that provide nutrient-rich habitats or reservoirs that can seed and fuel disease outbreaks once suitable environmental conditions are established (Mahmud et al. 2008). Given the complexity of marine ecosystems and their food webs, we have an incomplete understanding of how marine biodiversity more broadly contributes to pathogen persistence and dispersal in the marine environment.

Seagrass meadows are biodiversity hotspots, and deliver numerous ecosystem services in coastal regions. One such service that has been proposed, is reduction of pathogenic bacteria in water bodies surrounding meadows (Lamb et al. 2017). In the Baltic Sea, meadows of the dominant foundation species eelgrass (*Zostera marina*), have been associated with reduction of potentially pathogenic *Vibrio* spp. (Reusch et al. 2021). The mechanisms of such reductions remain unclear, but one proposed mechanism is increased sedimentation rates above seagrass meadows, which may contribute significantly to removal of *Vibrio* cells attached to particles in the water column, as vibrio often associate with surfaces (Huq et al. 1983, Datta et al. 2016, Kesy et al. 2020). In addition, several members of the eelgrass leaf microbiome feature antimicrobial activity against a range of pathogens (Tasdemir et al. 2024), suggesting that antagonistic interactions may play a role in suppressing *Vibrio* growth on or near seagrass surfaces. On the other hand, sequence variants classified as *Vibrio* spp. were also specifically detected in association with different seagrasses (Hassenrück et al. 2015, Bengtsson et al. 2017, Ugarelli et al. 2018, Sun et al. 2020, Tarquinio et al. 2021, Yan et al. 2021). Moreover, marine biofilms have in general been identified as reservoirs of pathogenic *Vibrio* (Shikuma & Hadfield 2010, Lutz et al. 2013), raising the possibility that also seagrass surfaces may harbor *Vibrio* strains capable of causing infections in humans and marine wildlife.

In this study, we aimed to provide broader insights about the ecology of *Vibrio* spp. in the context of seagrass meadow ecosystems. We investigated the prevalence of *Vibrio* spp., including the potentially pathogenic *V. vulnificus*, in and around seagrass meadows (*Z. marina*) around the island of Hiddensee (Mecklenburg-West Pomerania, Germany) in the south-eastern Baltic Sea. We used both culture-independent and culture-dependent tools to detect vibrios in water and sediment from vegetated and unvegetated areas, as well as on seagrass young and old leaf- and root surfaces, as previous studies have shown differences in microbiomes of (aging) plant organs (Ugarelli et al. 2018, Sanders-Smith et al. 2020). We hypothesized that (1) *Vibrio* spp. would be less prevalent in the water column above seagrass meadows, compared to unvegetated sites and that (2) surfaces in seagrass meadows such as leaves, roots and sediment particles are habitats for *Vibrio* spp., and can in this way function as reservoirs.

## 2. MATERIALS & METHODS

### 2.1 Study site and sampling

The study site is located around the Island of Hiddensee in the Baltic Sea, North-East Germany (Figure 1). Samples were collected in July 2019 from areas both with vegetation of the common eelgrass *Zostera marina* and from unvegetated areas. Geographical position, salinity, temperature, depth and secchi depth were documented at each site (Table 1).

**Figure 1:**
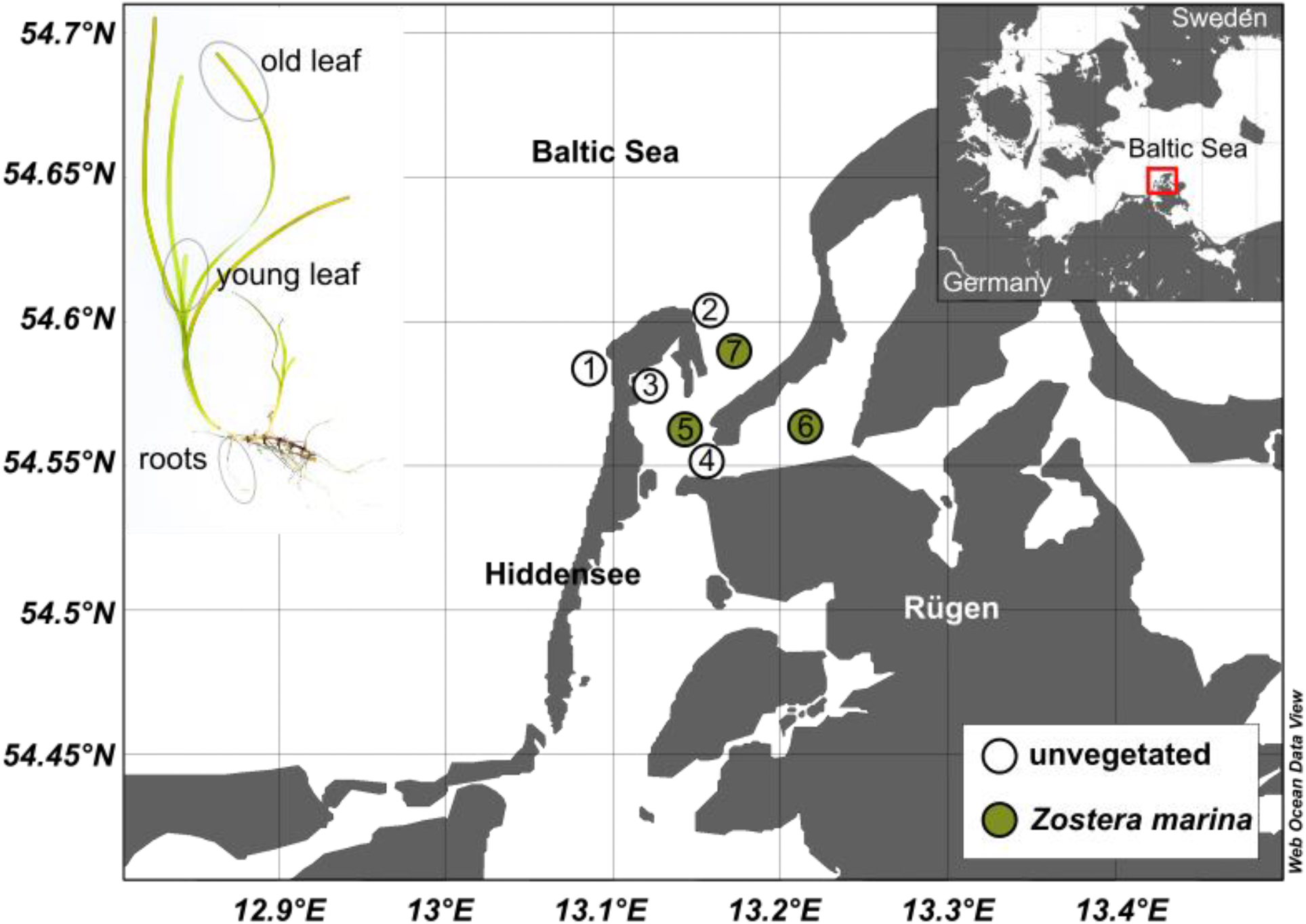
Sampling locations around the Island of Hiddensee with circles indicating site vegetation and overview of seagrass plant parts (left). Map of north-eastern Germany and the Baltic Sea (right) showing location of field site (red rectangle). The map was created using the web-based version of Ocean Data View (Schlitzer & Mieruch-Schnülle 2024).

**Table 1:** Geographical position and environmental conditions at sampling sites around the Island of Hiddensee, Germany.

| Site No. | Site | Latitude | Longitude | Salinity [psu] | Depth [m] | Secchi Depth [m] | Temperature [°C] |
| --- | --- | --- | --- | --- | --- | --- | --- |
| 1 | Kloster Beach | 54.58689 | 13.09703 | - | - | - | 19.5 |
| 2 | Lighthouse | 54.60093 | 13.1518 | 8.93 | 3 | >3 | 18.8 |
| 3 | Kloster Harbour | 54.5848 | 13.11122 | 9.49 | 1.9 | 1.7 | 20.8 |
| 4 | Vitter Loch | 54.55315 | 13.15817 | 9.26 | 5 | 2.2 | 20.2 |
| 5 | Kloster Loch | 54.5787 | 13.12525 | 9.35 | 2.6 | 2 | 20.8 |
| 6 | Rassower Strom | 54.56988 | 13.22657 | 9.31 | 3.4 | 1.7 | 20.6 |
| 7 | Libben | 54.58645 | 13.16232 | 9.24 | 3.7 | 3.7 | 18.2 |

**Table 2:** Results of PERMANOVA analysis (adonis2) to determine the role of sampling site (see Figure 1) and sample type (water, sediment, leaves and roots) on microbial community composition of all bacterial ASVs (16S), all eukaryotic ASVs (18S) and 16S ASVs classified as *Vibrio* spp. (*Vibrio* - 16S).

|  |  | df | SumSq | R<sup>2</sup> | F | p |
| --- | --- | --- | --- | --- | --- | --- |
| **16S** | *main* |  |  |  |  |  |
|  | site | 6 | 4.6483 | 0.26653 | 2.3014 | 0.001*** |
|  | sample type | 4 | 7.0234 | 0.40271 | 6.7424 | 0.001*** |
|  | *interaction* |  |  |  |  |  |
|  | site : sample type | 9 | 3.7273 | 0.21372 | 3.0304 | 0.001*** |
| **18S** | *main* |  |  |  |  |  |
|  | site | 6 | 4.5365 | 0.25 | 2.1111 | 0.001*** |
|  | sample type | 4 | 6.8844 | 0.3794 | 6.1134 | 0.001*** |
|  | *interaction* |  |  |  |  |  |
|  | site : sample type | 9 | 4.2170 | 0.23239 | 3.0937 | 0.001*** |
| ***Vibrio*-**<br>**16S** | *main* |  |  |  |  |  |
|  | site | 6 | 3.1286 | 0.21432 | 1.5458 | 0.006*** |
|  | sample type | 4 | 3.2391 | 0.22189 | 2.5664 | 0.001*** |
|  | *interaction* |  |  |  |  |  |
|  | site : sample type | 8 | 2.9579 | 0.20262 | 1.4416 | 0.010** |

**Table 3:** Results of a permutation test to determine the multivariate homogeneity of group dispersion (function permutest on output of function betadisper) depending on sampling site (see Figure 1) and sample type (water, sediment, leaves and roots) on microbial community composition of all bacterial ASVs (16S), all eukaryotic ASVs (18S) and 16S ASVs classified as *Vibrio* spp. (*Vibrio* - 16S).

|  |  | df | SumSq | MeanSumSq | F | p |
| --- | --- | --- | --- | --- | --- | --- |
| **16S** |  |  |  |  |  |  |
|  | site | 6 | 0.89364 | 0.148939 | 9.5736 | 0.001*** |
|  | sample type | 4 | 0.12155 | 0.0303863 | 4.2394 | 0.007*** |
| **18S** |  |  |  |  |  |  |
|  | site | 6 | 0.94774 | 0.157956 | 6.2488 | 0.001*** |
|  | sample type | 4 | 0.2337 | 0.0058432 | 0.3634 | 0.819 |
| ***Vibrio*-16S** |  |  |  |  |  |  |
|  | site | 6 | 0.85131 | 0.141884 | 8.5608 | 0.001*** |
|  | sample type | 4 | 0.21654 | 0.54135 | 5.7758 | 0.002** |

Whole plants of *Z. marina* were carefully harvested by hand via snorkeling and stored in zip-lock plastic bags. Sediment was collected in sterile 50 mL tubes either manually filled underwater or filled from a bulk sample retrieved with a Van Veen grab sampler from the boat. Water samples were collected by filling and sealing 200 mL screw cap flasks at a depth of 1 m underwater. All samples were stored cool and dark for 2-12 hours until further processing.

For analysis of microbial composition, using a culture-independent approach, subsamples consisting of either 10 cm long pieces of rinsed old and young seagrass leaves, rinsed roots or 2 mL of wet sediment were stored in 10 mL RNA later. For water samples, cells and particles from 200 mL of seawater were first collected on 0.22 µm Sterivex filters (polyethersulfone membrane) and preserved by filling the filter cartridge with RNA later, sealing the cartridges with parafilm and storing them individually in sterile 50 mL tubes. All samples were stored at 4 °C until DNA extraction.

For the cultivation-based approach, subsamples of either 2 cm of a rinsed and sterile cut leaf, 7 g sediment and 1 mL of seawater were added to separate sterile 50 mL tubes containing 10 mL sterile alkaline peptone water (10 g Peptone, 10 g NaCl in 100 mL H_2_O, pH 9) and subsequently stored at 4 °C during transport (<24h). For this approach, there was no differentiation between young and old leaves and no root samples were collected.

### 2.2 Enrichment cultivation of *Vibrio* spp. and 16S rRNA gene sequencing of colonies

Upon arrival to the lab, peptone water enrichments were incubated at room temperature (27 °C) for 24 h and then each of them were cryopreserved by adding 50 % glycerol and stored at −80 °C. A serial dilution of each peptone water enrichment was spread evenly on separate TCBS agar plates, incubated at 30 °C and colony forming units (CFU) counted after 48 h. 200 colonies selected according to color were picked and stored in 100 µL nuclease-free water at −20 °C until processing for colony-PCR using the primer pair B341F (5’-CCTACGGGNGGCWGCAG-3’) and B806R (5’-GGACTACHVGGGTATCTAAT-3’) (Klindworth et al. 2013) for amplification of the hypervariable V3-V4 regions of the 16S rRNA gene. After agarose gel electrophoresis, fragment clean-up (ZR 96 DNA clean up kit, Zymo Research) and quantification via UV-absorbance using a Nanodrop system (Thermo Fisher), 5 ng µL^-1^ of the final PCR-products were sent for Sanger sequencing (Eurofins Genomics, Berlin). The colony sequences have been submitted to Genbank under accession numbers PP806658 - PP806832.

The number of colony forming units (CFU) from the enrichment cultivation was calculated as CFU per mL peptone water. To account for a bias in selectivity of the TCBS agar, a factor correcting for potentially false-identification of *Vibrio* spp. based on the sequencing results of picked colonies was included in the calculation for each sample type and both sampling days individually (Supplementary Table S2). A Kruskal-Wallis-Test was performed to test if CFU concentration in the water samples differed significantly between vegetated and unvegetated sites.

### 2.3 DNA extraction, PCR amplification and sequencing

To remove the excess RNA later, leaves were rinsed with 1400 µL pre-cooled PBS and cut into smaller pieces, 1 g of sediment was pelleted via centrifugation, the supernatant containing RNA later removed, the remaining sediment washed with 1400 µL PBS and 0.25 g of sediment was transferred to a tube. The Sterivex filter cartridges were cracked open with a hammer, the membrane removed with sterile forceps, rinsed with 1400 µL PBS, cut with sterile blades and used for further extraction.

DNA was isolated using the DNeasy PowerSoil Pro kit (Qiagen) according to the instructions with prior sonication (2 x 7 min on ice) and bead beating (30 s at 4 m s^-1^ + 45 s at 5 m s^-1^, with leaf samples only exposed to the 1^st^ step). Slightly different cell dislodgement methods (swabbing versus bead beating) were tested to increase DNA yield (see Supplementary material for a detailed description), however, a PERMANOVA including the dislodgement method showed that these had no significant influence in explaining community composition (PERMANOVA test for 16S data: R^2^ = 0.059, p = 0.095; 18S data: R^2^= 0.07, p = 0.097; see Supplement). DNA concentration was measured using a Qubit fluorometer (Thermo Fisher) and aliquots of 7 ng µL^-1^ from the crude extracts were shipped to LGC Genomics who performed the amplicon library preparation and paired-end sequencing (2 x 300 bp) on the Illumina Mi-Seq-platform with the V3 kit (Illumina Inc., Berlin, Germany). For analyzing bacterial composition and *Vibrio* abundance, primers targeting the V4 region of the 16S rRNA gene were used (515f: 5‘-GTGYCAGCMGCCGCGGTAA-3‘, 806r: 5‘-GGACTACNVGGGTWTCTAAT-3’ (Walters et al. 2016)) and primers targeting the V7 region of the 18S rRNA gene were used to identify potential hosts for vibrios (F-1183mod: 5’-AATTTGACTCAACRCGGG-3’, R-1443mod: 5’-GRGCATCACAGACCTG-3’ (Ray et al. 2016)). The amplicon sequence data has been submitted to the European Nucleotide Archive (ENA) under the project number PRJEB75525.

### 2.4 Absolute quantification with ddPCR

Molecular quantification of *Vibrio vulnificus* was performed on the same DNA extracts that were used for sequencing utilizing Bio-Rad’s X200 Droplet Digital PCR systems and the primer pair vvh785F (TTCCAACTTCAAACCGAACTATGAC) and vvh990R (ATTCCAGTCGATGCGAATACGTTG) which has been proven to specifically target the *vvh* gene that encodes a *V. vulnificus*-specific hemolysin (Panicker et al. 2004). A 22 µl ddPCR reaction mix consisted of 11 µL EvaGreen digital PCR Supermix, 100 nM forward/ reverse primer and 0.1-1 ng template DNA. Samples were measured in a 10-fold dilution series. The positive reaction included template DNA from *V. vulnificus* (DSMZ 10143), MilliQ-water was used as a negative control. Samples (20 µL) mixed with Droplet Generation Oil (70 µL) were partitioned into nano-sized droplets using the X200 Droplet Generator, manually transferred to a 96-well plate and heat-sealed with a foil cover. Amplification was performed using a thermocycler with initial denaturation for 5 min at 95 °C, followed by 40 cycles of denaturation (95 °C, 30 s), primer annealing (58.5 °C, 1 min) and target extension (72 °C, 2 min) with a ramp-rate of 2 °C s^-1^ for each step. After amplification, samples were read with the Bio-Rad Droplet Reader.

Absolute quantification of target gene copies was performed using Quanta Soft version 1.7.4.0917 (Bio-Rad) with default ABS settings and reactions with <10.000 droplets were excluded from further analysis. A threshold to separate positive and negative droplets was set manually after visual inspection of positive and negative controls. The amount of copies ng^-1^ DNA was calculated by multiplying the reaction copies µL^-1^ provided by the Software with the reaction volume (22 µL) divided by the volume of DNA used for the reaction (µL), dividing the result by the concentration (ng µL^-1^) of total DNA sample.

### 2.5 Data processing and bioinformatics

The demultiplexed and clipped sequences obtained from LGC Genomics were denoised, filtered for chimeric sequences and grouped into amplicon sequence variants (ASV) using the dada2 pipeline (Callahan et al. 2016). Taxonomic annotation of ASVs was performed using the rdp classifier with SILVA version 138 (16S rRNA gene) and SILVA version 132 (18S rRNA gene) as reference databases (Pruesse et al. 2007). The generated ASV tables were filtered either to remove organisms other than bacteria and archaea for the 16S rRNA dataset (e.g. eukaryotic sequences such as from mitochondria, chloroplasts) or sequences originating from *Z. marina* in the eukaryotic 18S rRNA dataset. Prior to analysis, ASV data were normalized to relative abundances.

### 2.6 Differential abundance analysis

To test if seagrass presence had an influence on *Vibrio* abundances and other community members in the water samples, a differential abundance analysis was performed on the read-based dataset using the DESeq2 package at the genus level data (Love et al. 2014). We also tested if seagrass tissue type (young leaves, old leaves, roots) differed in their *Vibrio* abundances. For that, samples were split into a water and a seagrass dataset. To reduce spurious signals in the differential abundance analysis, genera were filtered for a prevalence of 50 %, so that only genera occurring in at least 50 % of the samples were retained for the analysis. This was done for each dataset individually. Differentially abundant genera were identified based on their log2 fold change in read abundance using the DESeq function with the default testing framework (Wald test and parametric dispersion fitting of mean intensities) and the above described contrasts. As there was only one sediment sample from unvegetated sites, no such test was performed for sediment samples.

### 2.7 Community composition analysis

Analyses of community composition were performed by processing ASV tables in the R environment (R Core Team 2023) with the phyloseq (McMurdie & Holmes 2013), vegan (Oksanen et al. 2022) and ggplot2 (Wickham 2009) packages. To compare microbial community composition among the sites, compositional dissimilarity matrices were calculated from Bray-Curtis-distances using the distance function of the phyloseq package. The betadisper function of the vegan package was used to quantify the compositional variance within the sample types. This was done by estimating the average distance of individual group members to the group centroid in multivariate space based on previously calculated dissimilarity matrices and visualized using principal coordinates analysis (PCoA).

To determine the influence of sampling site and sample type on microbial community composition, we used the vegan package (Oksanen et al. 2022) to perform a PERMANOVA test using the adonis2 function with 999 permutations for all bacterial ASVs, eukaryotic ASVs and ASVs classified as *Vibrio* spp. We also included a permutation (permutest of the vegan package) to analyze the multivariate homogeneity of group dispersion for microbial community composition (betadisper results) depending on sampling site and sample type.

### 2.8 Correlation between 16S, 18S and *Vibrio* data

To investigate the influence of biotic factors on *Vibrio* community composition, a Procrustes analysis was performed using the Procrustes and protest function in vegan (Oksanen et al. 2022). To detect correlations with prokaryotic and/ or eukaryotic communities, Procrustes was applied to Principal Coordinates Analysis ordinations (PCoA, based on Bray-Curtis dissimilarities) of the 16S and 18S rRNA community (all *Vibrio* ASVs excluded) and of all *Vibrio* ASVs. The same procedure was used for the 18S rRNA dataset. To separate the influence of sample type and sample site from any biotic influence, a third distance matrix encoding the environmental parameters ‘sample type’ and ‘site’ was created and a partial Mantel test (vegan) performed.

### 2.9 Construction of a phylogenetic tree for sequence comparison

To compare the partial 16S *Vibrio* sequences obtained from the cultivation approach with those found in the amplicon dataset, a phylogenetic tree was constructed using ARB6 (Ludwig et al. 2004) and the SILVA 138.1 SSURef NR99 reference database (Yilmaz et al. 2014). Sequences were first aligned using the AlignSeqs function from the DECIPHER package in R (Wright 2016) and then trimmed to the same length within MEGA11 (Tamura et al. 2021), resulting in ∼150 bp amplicon reads. Those were transferred into ARB format using the online SINA aligner (Pruesse et al. 2022) and merged with the SILVA 138.1 SSURef NR99 reference database within ARB. To create a scaffold tree with related *Vibrio* type strains, 336 *Vibrio* type strains were selected, and a phylogenetic tree was constructed using RAxML via the online ACT-tool (Pruesse et al. 2012, Stamatakis 2014). The tree was rooted using *Thioalkalispiraceae* sequences (n = 74) as an outgroup. Environmental *Vibrio* sequences were then added to this scaffold tree via the quick-add command in ARB. The tree was refined by successively removing type strains on distant branches. Finally, identical environmental *Vibrio* sequences from the same approach were collapsed into groups for visualization.

## 3. RESULTS

Members of the genus *Vibrio* were detected from several of the samples with all three analytical approaches used in this study. The relative abundance of ASVs classified as *Vibrio* spp. in the 16S rRNA dataset varied substantially with sample type and ranged between 0-8 % of the total bacterial community (Figure 2a). The highest relative abundances were detected on *Z. marina* roots, where *Vibrio* spp. ASVs were significantly overrepresented compared to both old and young leaves (Supplementary tables S3, S4). Based on the 16S rRNA gene amplicon data, there were no differentially abundant genera (including *Vibrio*) in water samples collected above *Z. marina* meadows compared to unvegetated areas.

**Figure 2:**
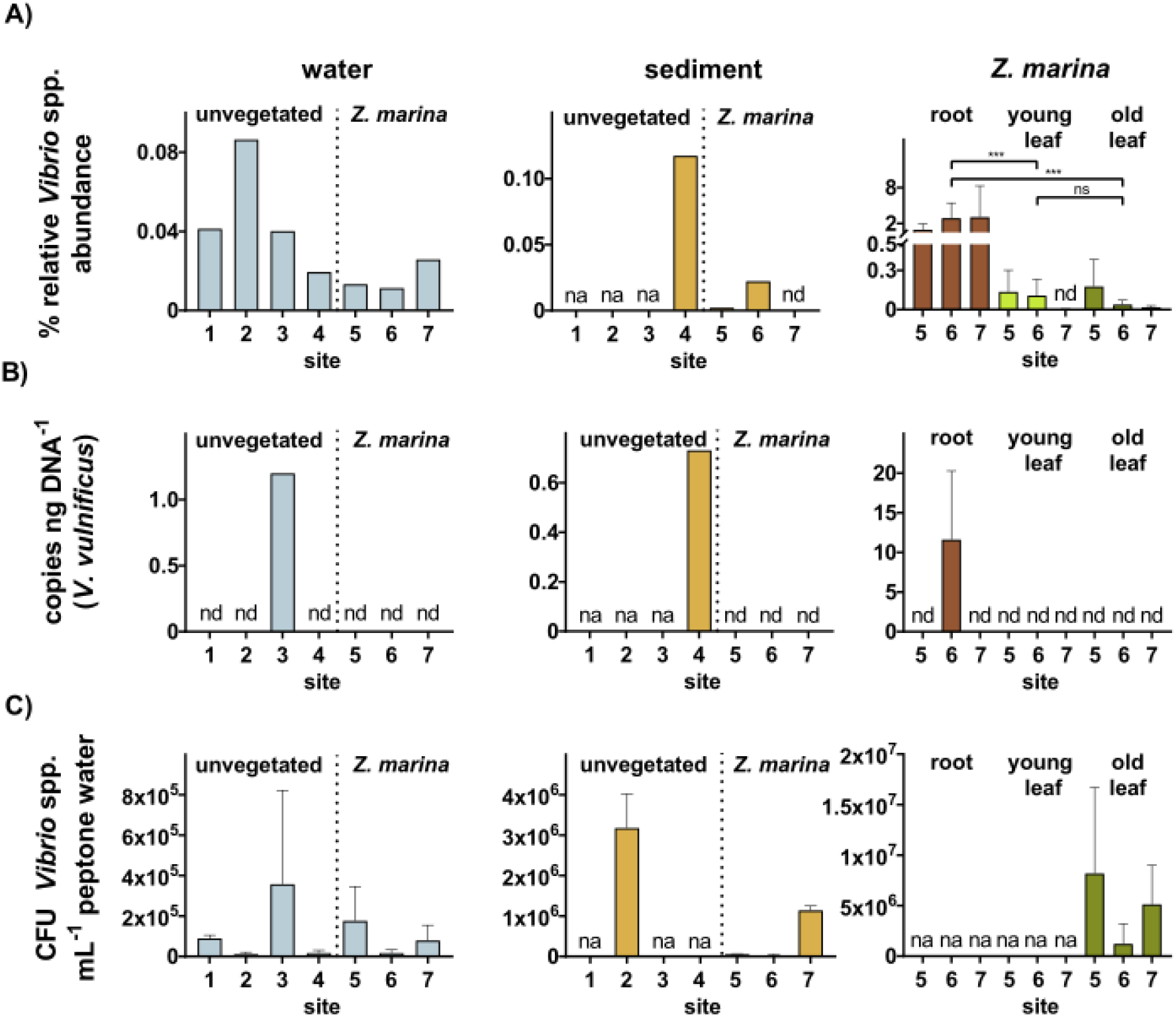
Presence of *Vibrio* spp. in water, sediment and on plant parts (young and old leaves, roots) from unvegetated and vegetated sites as detected via three independent methodologies **A)** relative abundance from amplified 16S rRNA genes taxonomically classified as *Vibrio* spp., **B)** absolute abundance of *Vibrio vulnificus* from ddPCR analysis and **C)** Colony-forming units of *Vibrio* spp. from a selective culturing approach (Peptone water enrichment followed by TCBS agar, compensation for false negative via sequencing). No root samples were taken for the cultivation approach. Differential abundance analysis on plant parts revealed significant differences of *Vibrio* spp. on roots compared to old and young leaves (A, asterisks), the difference between young and old leaves was not significant (ns), as was the difference between water samples from unvegetated and *Z. marina* sites. na = sample not available, nd = not detected.

Using ddPCR with specific primers, we also detected the potentially pathogenic *V. vulnificus* in sediment and water from unvegetated areas and also associated to *Z. marina* roots at one of the sampling sites (Figure 2b). Statistical comparison between sites or sample types was not possible due to low prevalence.

The cultivation-based approach, using selective TCBS agar and compensation for false negative identification via sequencing of colonies (Supplementary Table S2), revealed an abundance of *Vibrio* spp. colony forming units, especially on *Z. marina* leaves, but also in water samples (Figure 2c). There was no significant difference in the amount of CFUs detected in water from unvegetated sited compared to seagrass meadows (p = 0.7237). Root samples were not collected for *Vibrio* cultivation, so the abundance of *Vibrio* CFUs on *Z. marina* roots could not be determined.

Among the colonies picked for partial 16S rRNA gene sequencing (n = 200), 81 % belonged to the genus *Vibrio*, while 13.5 % belonged to the genera *Shewanella*, *Photobacterium*, *Morganella* and *Aeromonas*. The remaining 5.5 % could not be taxonomically classified due to poor sequence quality. A phylogenetic analysis revealed that the cultivation-based approach (colony-derived sequences) and the 16S amplicon sequencing (ASVs) detected very similar or identical *Vibrio* sequence variants (Figure 3). The detected *Vibrio* sequence variants were related to sequences from several cultured *Vibrio* reference strains available in public databases.

**Figure 3:**
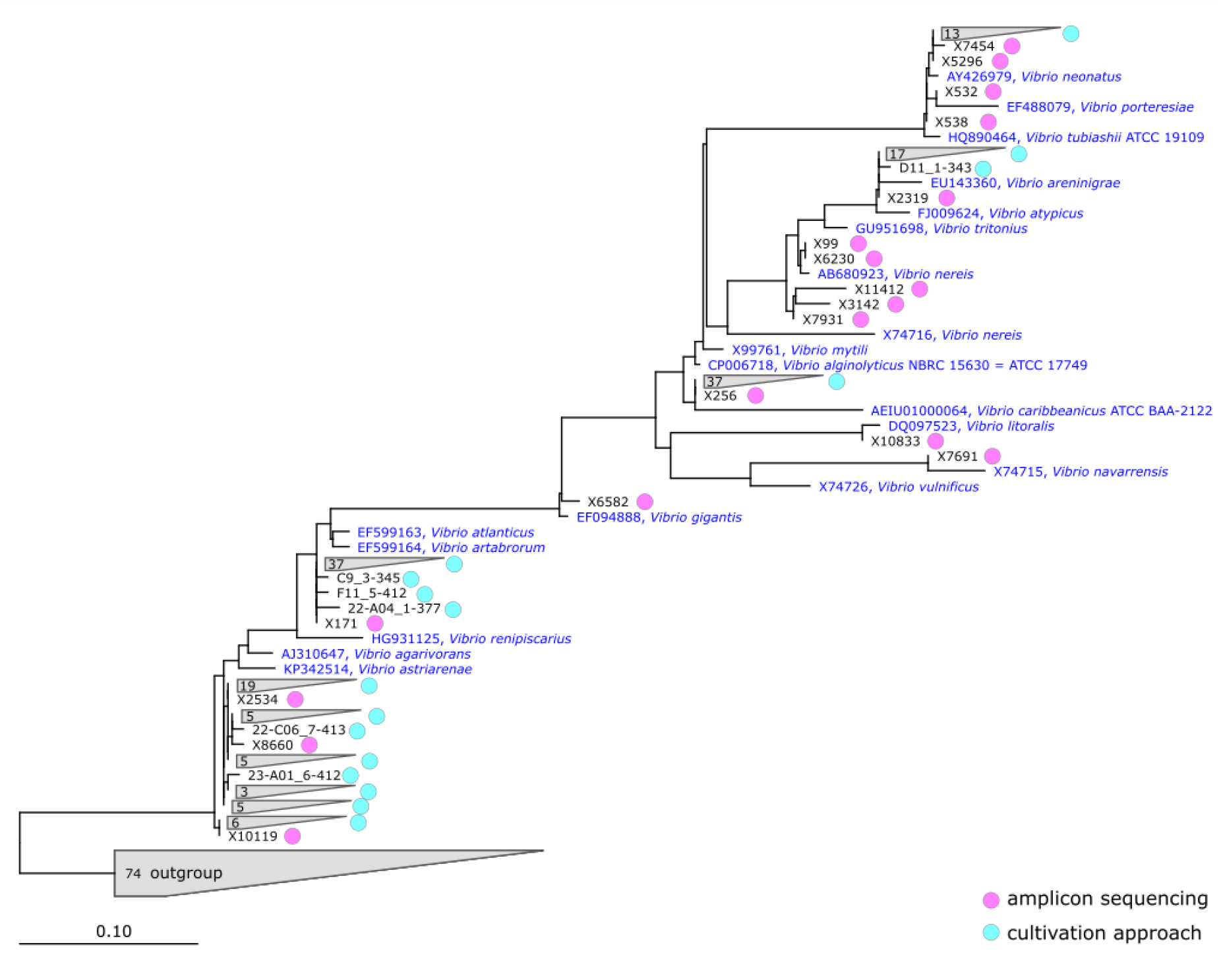
Phylogenetic tree of colonies- and ASVs partial 16S rRNA gene sequences. The tree was constructed using ARB6, RAxML, and the SILVA 138.1 SSURef NR99. Black names indicate sequences generated in this study, blue names are type strains. 100 % identical sequences from colonies (cultivation approach) are aggregated in grey triangles, numbers indicate amount of sequences within aggregates. Outgroup sequences belonged to *Thioalkalispiraceae*. Bar: 10 substitutions per 100 nucleotides.

However, identification to species level was not possible as the low taxonomic resolution of the partial 16S rRNA gene sequences generated in this study is of limited value for this purpose in vibrios (Thompson et al. 2009).

Amplicon sequencing of the 16S rRNA gene revealed that the overall bacterial community composition was highly dependent on sample type, explaining 40 % of variation (R^2^ = 0.40, p<0.001) according to PERMANOVA analysis (Table 1), while sampling site played a relatively minor but significant role (R^2^ = 0.27, p<0.001). A PCoA ordination showed that water samples were strongly separated from all other sample types, while young and old leaves, as well as roots and sediments, were more similar (Figures 4a, 5a). Analogously, eukaryotic community composition indicating the occurrence of possible microeukaryotic predators or hosts of *Vibrio* spp., was also mainly explained by sample type (R^2^ = 0.38, p<0.001) compared to sample site (R^2^ = 0.25, p<0.001). Seagrass leaves displayed a distinct eukaryotic community compared to all other sample types (Figures 4b, 5b). Unlike the overall microbial community (bacteria and eukaryotes), the composition of *Vibrio* spp. based on 16S rRNA gene amplicon sequencing was similarly explained by both sample type (R^2^ = 0.21, p<0.006) and sampling site (R^2^ = 0.22, p<0.001). Only root samples clustered separately from other sample types in the PCoA ordination (Figure 4c).

**Figure 4:**
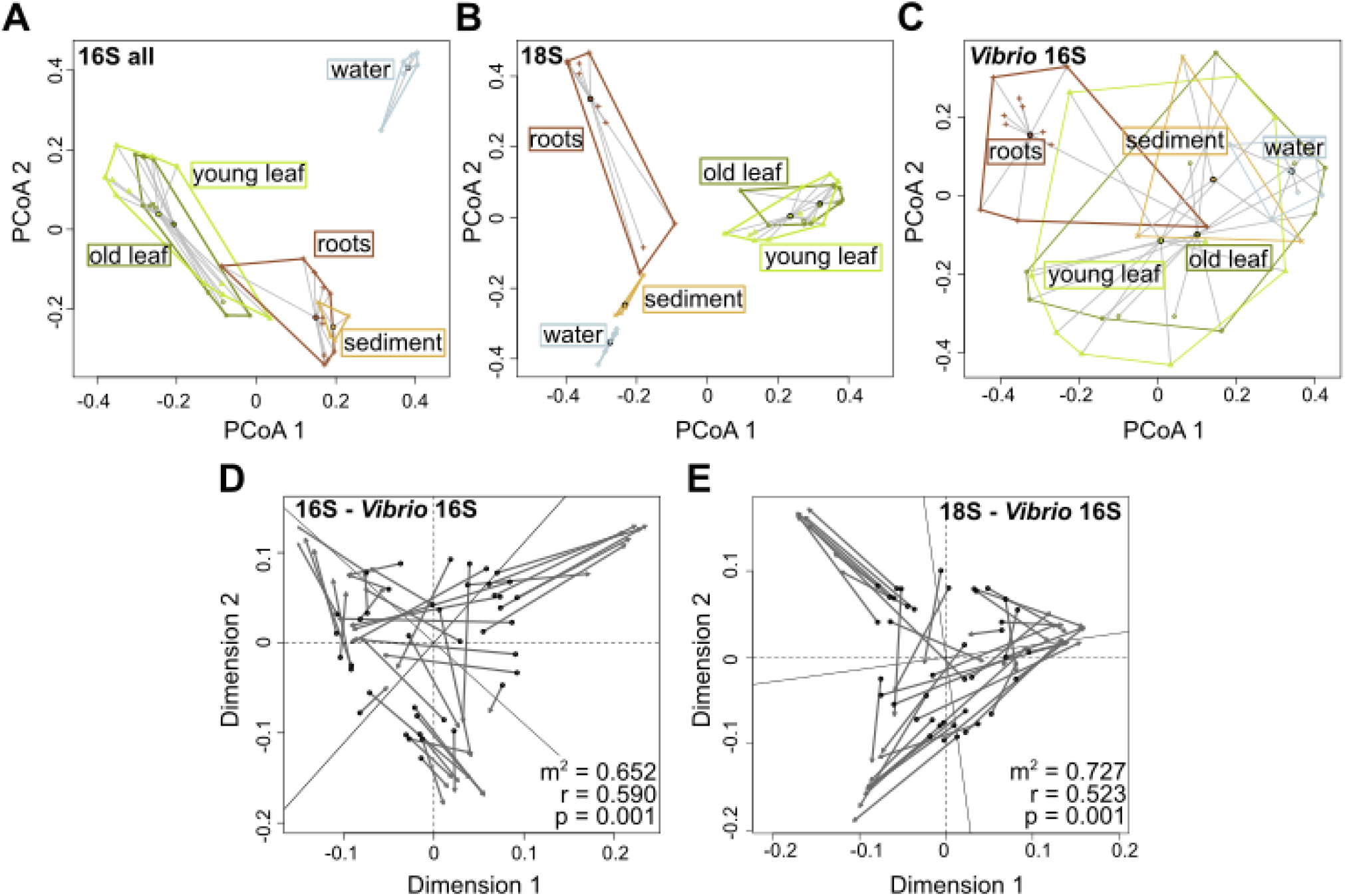
Composition and comparison of prokaryotic, eukaryotic and *Vibrio spp.* relative abundances. Principal Coordinates Analysis (PCoA) plots showing variation between sample types based on Bray-Curtis distances of **A)** prokaryotic and **B)** eukaryotic communities as well as **C)** variation of *Vibrio* spp. Scatter plots of generalized Procrustes analysis mapping **D)** prokaryotic phylogenetic composition and **E)** eukaryotic phylogenetic composition to sample ordinations based on relative abundance of *Vibrio* spp. from the same set of samples. Longer lines between a sample community eigenvalue (points) and its concurrent *Vibrio* spp. eigenvalue (arrows) indicate greater discordance between datasets for that sample. In all comparisons the m^2^-value was considered significant (p<0.001).

A Procrustes analysis was performed to assess the influence of the biotic environment (i.e. the microbial communities coexisting with *Vibrio* spp.) on *Vibrio* composition in seagrass meadows. A correlation of *Vibrio* spp. ASV distribution with the overall bacterial as well as the overall eukaryotic community composition revealed, that both bacterial and eukaryotic community composition correlated significantly with *Vibrio* spp. composition, but this correlation was somewhat stronger for bacterial (r = 0.59, p = 0.001) than for eukaryote communities (r = 0.52, p = 0.001). However, after partitioning out the strong variation caused by sample type and sample site using a partial Mantel test, the eukaryotic community had no significant influence (eukaryotic community: r = 0.039, p = 0.183, prokaryotic community: r = 0.178, p<0.001).

## 4. DISCUSSION

In this study, we investigated the abundance and diversity of *Vibrio* spp. in a specific ecological setting, namely seagrass meadow ecosystems. Our findings confirm that *Vibrio* spp. are indeed present in several compartments of seagrass meadows and are represented by several different taxa, including in some cases the potentially pathogenic *Vibrio vulnificus*. Our study adds to a growing body of literature trying to link pathogen abundance and distribution to the dynamics and features of ecosystems that are facing the consequences of global change (Sterk et al. 2013, Vezzulli et al. 2013, Cohen et al. 2018). In order to understand why pathogens such as *V. vulnificus* sometimes depart from the marine environment and infect for example humans, it is essential to understand how these bacteria interact and respond to the state of their environment. Here, we shed light on some aspects of *Vibrio* spp. ecology in and around seagrass meadows in the south-eastern Baltic Sea, an area impacted by several environmental stressors such as rising seawater temperatures and eutrophication (Reusch et al. 2018).

### No difference in *Vibrio* abundance in the water column above seagrass beds compared to unvegetated sites

We hypothesized that *Vibrio* spp. would be less prevalent in the water column above seagrass meadows as compared to unvegetated sites. This was based on the results from two recent studies investigating either several types of pathogenic bacteria (Lamb et al. 2017), or pathogenic vibrios specifically (Reusch et al. 2021) in seagrass meadows. However, we could not detect such a difference with any of the three analytical approaches used. It is still possible that actual differences would be hidden by substantial spatial variation in their distribution, not least since our sampling design only included three sites with seagrass meadows and four unvegetated sites. Our study area is located in a complex coastline setting characterized by large bays and lagoons with various degrees of wave exposure and water exchange (Kniebusch et al. 2019). This adds to the natural variability of e.g. temperature and salinity regimes which are known to influence the growth and environmental prevalence of some *Vibrio* species (Eiler et al. 2006), and could thus mask an effect of seagrass presence. Unlike another recent study from the Baltic Sea (Reusch et al. 2021), we did not assess differences in *Vibrio* abundances on a small spatial scale (over a few meters) in and around individual meadows, but instead sampled sites that were typically several kilometers apart resulting in no information on local environmental variation, which might be of importance. Therefore, despite our findings, we cannot conclusively exclude the possibility that seagrass meadows in our study area function as natural filters that remove vibrios and other pathogenic bacteria from the water column, e.g. by attaching bacteria to plant material. In line with this, the highest *Vibrio* spp. (relative) abundances in our study were observed for sites without seagrass vegetation with all three methodological approaches, but those differences were not significant.

Currently, there is no universal methodological approach which can be used to compare results between existing studies on abundance of vibrios in seagrass ecosystems and the choice of methodology may have a large impact on results. For example, we chose an enrichment cultivation approach including both peptone water and TCBS agar (Lesmana et al. 1985) which may select for certain populations of vibrios yet excluded others compared to other enrichment media, e.g. CHROMagar (Reusch et al. 2021). Sequencing of selected colonies also showed that selectivity of methodologies could still be improved and reliable identification requires a combination of techniques. Combined TCBS agar and CHROMagar have been evaluated and used for detection and identification of vibrios in the Baltic Sea (Glackin et al. 2024) and may be a promising low-cost quantification method. The use of ddPCR and specific *Vibrio* primers has emerged as a reliable cultivation-independent quantification method (Möller et al. 2021). Until a unified methodology is adopted, validation and comparison between studies will remain difficult.

### *Vibrio* spp. are abundant on *Z. marina* roots

Seagrass roots stood out as the habitat exhibiting the highest relative abundances of *Vibrio* spp. ASVs in our 16S rRNA amplicon sequencing survey. *V. vulnificus* was also detected in higher absolute abundance on root samples from one of the sites (site 6, Rassower Strom) relative to other samples. Our observations are supported by results from previous studies which also observed *Vibrio* spp. in association with seagrass roots: High proportions of *Vibrio* spp. were detected in an investigation of rhizosphere sediments of *Z. marina* and *Z. japonica* in Swan lake in China (Sun et al. 2020). The authors noted that vibrios were overrepresented in seagrass habitats compared to degraded and bare sediments. This led them to speculate that the high proportions may be linked to co-occurring macrobenthic species, such as Bivalves. In 2020, Wang et al. found *Vibrio* to be an indicator taxon for roots in fertilized seagrass and much earlier nitrogen-fixing *Vibrio* strains were also isolated from Z. *marina* roots (Shieh et al. 1989). Thus, vibrios may not be the most abundant members of the seagrass root microbiome, but they nonetheless seem to be over-represented in root-associated communities as compared to other compartments (water, sediment, leaves, Figure 4).

Many vibrios are known to associate with larger organisms (Huq et al. 1983), form biofilms on biotic and abiotic substrates (Alam et al 2007). This has been shown for a diverse range of surfaces, such as wood, chitin, and plastics (Shikuma & Hadfield 2010, Datta et al. 2016, Oberbeckmann et al. 2017, Kesy et al. 2020). Vibrios are generally considered to feature a feast- and famine growth-strategy (Heidelberg et al. 2000, Thingstad et al. 2022), being able to quickly colonize newly available habitats and exploit episodic carbon and nutrient pulses (Westrich et al. 2016). The leaking of organic compounds through the seagrass roots, which follows diurnal cycles but is also locally variable, may provide such a habitat (Wood & Hayasaka 1981, Donnelly & Herbert 1998, Nielsen et al. 2001, Kurtz et al. 2003, Brodersen et al. 2018, Rotini et al. 2020).

### *Vibrio* spp. communities are influenced by both habitat type and site, and correlate with other microbial community structure

The strong influence of sample type and to a lesser extend also sampling site on microbial community composition was expected, as similar observations have been made in previous studies (Bengtsson et al. 2017, Fahimipour et al. 2017). The clear separation of microbial community types across different plant parts, as well as between the plant and water and sediment habitats clearly confirm that the seagrass host is a highly selective environment which features specialized microbiomes (Ugarelli et al. 2018, Tarquinio et al. 2019). The composition of *Vibrio* populations was influenced by the same structuring factors as the overall bacterial community, but with sample type (leaves, roots, sediment) having a relatively lower influence. This may indicate that other environmental factors such as temperature and salinity (Baker-Austin et al. 2013, Vezzulli et al. 2013, Baker-Austin et al. 2018), that are not directly related to the seagrasses, may play a larger role in selecting for specific vibrios compared to other members of the broader bacterial community.

We also tested for correlations of *Vibrio* communities with non-*Vibrio* bacteria and (microbial) eukaryotes in order to have an indication of whether biotic interactions between microbes may play a role in selecting for *Vibrio* ASVs that colonize seagrass surfaces, sediments, and the water column. The relatively strong correlation between *Vibrio* composition and bacterial community composition, even after partitioning out the influence of the sampling site, could either suggest that a broader set of organisms respond to the same environmental variables or that biotic interactions with other bacteria, such as competition, play a role in determining which vibrios persist in these environments (Tasdemir et al. 2024). As fast-growing, relatively large-celled marine bacteria, vibrios may outcompete other bacteria when resources are abundant, but at the same time be more vulnerable to competition and predation during periods when such resources are in short supply (Thingstad et al. 2022).

Even though *Vibrio* composition did not significantly correlate with microbial eukaryote composition after partitioning out parameters with a partial Mantel test, microeukaryotic bacterivory mediated by ciliates or bryozoans may still be relevant for structuring *Vibrio* communities in these habitats as these microeukaryotes were particularly abundant on seagrass leaves (Figure 5) and specific predators may interact with specific vibrio populations. The presence of eukaryote bacterivores utilizing different feeding modes selecting for prey in combination with lower availability of organic carbon may represent conditions that limit the abundance and proliferation of vibrios in seagrass meadow ecosystems (Macek et al. 1997, Beardsley et al. 2003, Pernthaler 2005).

**Figure 5:**
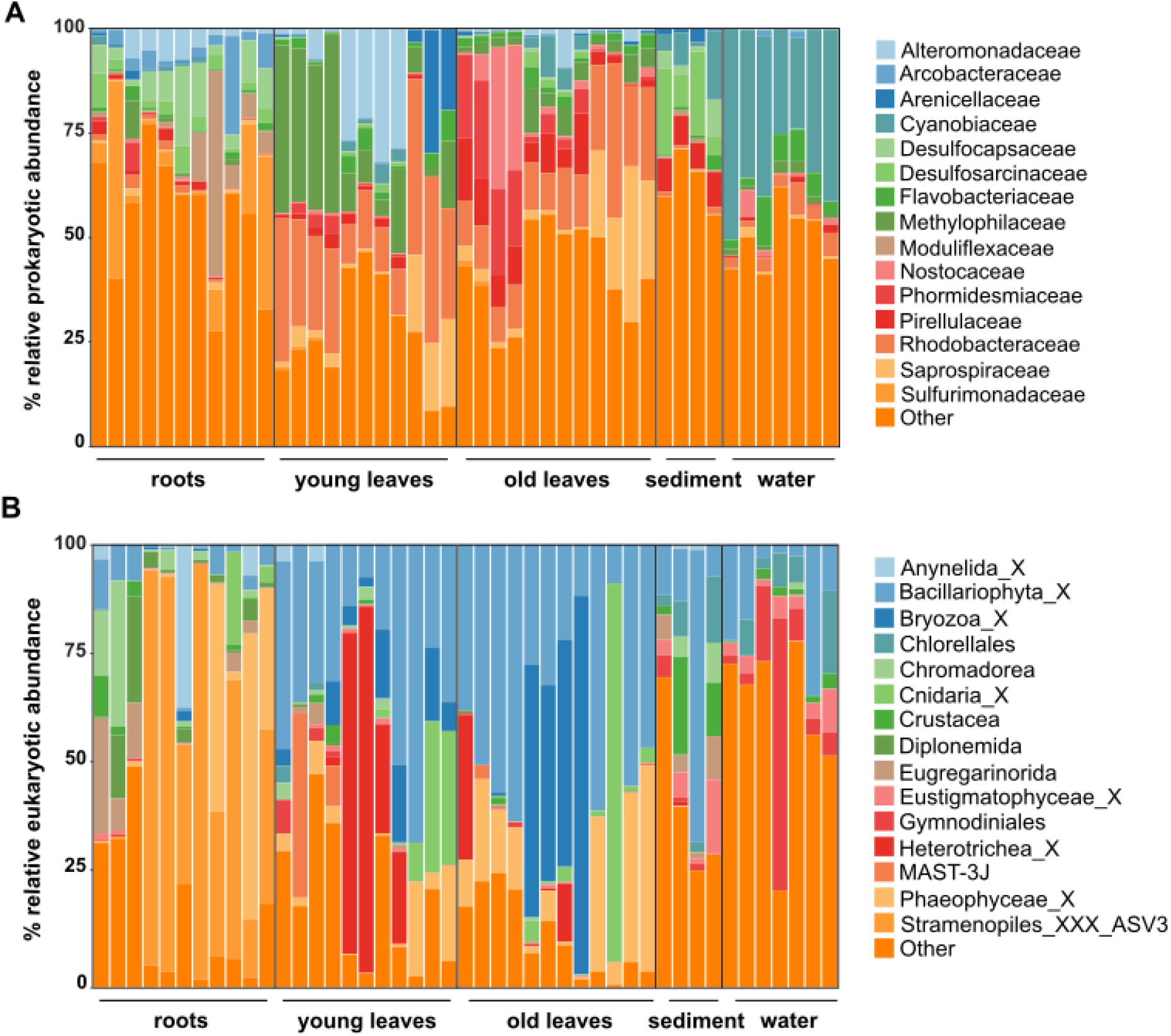
Prokaryotic (A) and eukaryotic (B) community composition across roots, young leaves and old leaves of *Z. marina* as well as surrounding sediment and water. Unique ASVs were summarized at family level and the representation of taxonomic groups within each sample are plotted. The overview is limited to the most abundant 15 families across all sample types, the remaining taxa are summarized as ‘Other’.

Correlative analyses such as Procrustes and partial Mantel tests employed to associate overall bacterial and eukaryotic communities with *Vibrio* community composition have to be interpreted with caution. Rather than interactions between organisms, correlations may reflect underlying correlations with unmeasured environmental factors, despite partitioning out the effect of measured factors.

### Implications and future perspectives

Our findings could neither conclusively confirm or refute the emerging idea that seagrass meadows function as filters that remove pathogenic bacteria from the water column (Lamb et al. 2017, Webb et al. 2019, Reusch et al. 2021). In order to clarify whether this potentially important ecosystem service is relevant in coastal ecosystems, additional studies need to be carried out in different coastal areas, but also on different spatial scales to account for the dynamic nature of coastal habitats. Another aspect that remains enigmatic, is the mechanism by which such a reduction in pathogen load could take place. One suggested mechanism is accelerated sedimentation above seagrass meadows, a process that may lead to sediment burial of mainly particle-associated pathogenic bacteria in seagrass meadow sediments (Lamb et al. 2017, Reusch et al. 2021). In addition, antimicrobial compounds produced by the seagrass (Guan et al. 2017), but also by its associated microbiome (Tasdemir et al. 2024) have been proposed to suppress growth of potentially pathogenic bacteria. A less investigated but potentially important removal mechanism is predation by epibiotic microbial eukaryotes, such as the abundant filter-feeding bryozoans, hydrozoans, rotifers and ciliates that colonize seagrass leaves. Such predators may preferentially remove vibrios from the water column and be an important sink (Worden et al. 2006).

Similarly, raptorial predators such as ciliates, flagellates and amoeba inhabiting seagrass leaf-surface biofilms may ingest *Vibrio* cells settling on surfaces. These interactions may not act on entire genera but between specific predators and specific vibrio populations.

We identified seagrass roots as habitats enriched in vibrios, a finding that is supported by previous studies (Martin et al. 2020, Sun et al. 2020, Wang et al. 2020, Yan et al. 2021). So far, the main focus of studies to assess the influence of root-associated microorganisms on seagrass health have mostly been on organisms mediating sulfur-cycling, coupled with nitrogen-fixation, a metabolic mode that seems to be dominant in the seagrass rhizosphere (Ugarelli et al. 2018, Tarquinio et al. 2019). We found that *Vibrio* spp. constitute a considerable fraction of the rhizosphere microbiome and is consistently present. Yan et al. hypothesized 2021 that vibrios could be an indicator of poor seagrass health. In contrast, in 2020 Martin et al. identified an increase in sulfur-cycling microbial lineages as a potential indicator of poor plant health while vibrios were considered to be part of the core root microbiome. In the present study, we did not measure seagrass health indices, however, the sampled seagrass sites harbor a persistent seagrass meadow, indicating that seagrasses here are at least healthy enough to survive and persist (pers. observation).

Many members of the genus *Vibrio* are known to be opportunistic pathogens, yet given that vibrios isolated from seagrass roots seem to be diazotrophs (Shieh et al. 1989, Jose et al. 2014, Garcias-Bonet et al. 2016), they could potentially also be beneficial for the plant rather than just being an opportunistic colonizer. Given their widespread presence in seagrass meadows and the importance of such habitats for the functioning of the coastal ecosystem, the ecological role of vibrios in such habitats should be further studied.

## Acknowledgements

We thank Dr. Sven Dahlke and Dr. Irmgard Blindow from the Biological Station on Hiddensee for essential assistance and infrastructure support during sampling. The National Park “Vorpommersche Boddenlandschaft” provided permission to access the sampling sites. We also thank Heike Benterbusch from IOW for assistance with ddPCR. This study was funded by the German Federal Ministry of Education and Research via the project SeaStore (BMBF 03F0859C, PI M.M. Bengtsson), as well as by EU Biodiversa+ via project BaltVib (16LC2022A, PI M. Labrenz). Additional support was provided through a PhD stipend awarded to R. Gebbe by the state of Mecklenburg-Vorpommern (Landesgraduiertenstipend) and a PhD stipend awarded to A. Brauer by the German Federal Environmental Foundation (DBU). The work was part of D. Hallieŕs Diploma-Thesis.

## Author contributions

MMB designed and supervised the study with contributions from SB and ML. MMB, AB and DH executed sampling and data generation. RG and KK analyzed the data with contributions from AB and DH. RG, KK and MMB interpreted the results, wrote the first draft of the manuscript and RG visualized the data. All co-authors provided their input on the manuscript and helped editing. All co-authors approved the submitted manuscript.

## Notes

### Competing Interest Statement

The authors have declared no competing interest.

