## Supplementary table S1, Supplementary table S2, Supplementary table S3, Supplementary table S4 for "The ecology of potentially pathogenic *Vibrio* spp. in a seagrass meadow ecosystem"

**Cell dislodgment pre-trials**

Three different methods for detaching the biofilm from sample surfaces were used: swabbing (designation „method S“), bead beating (designation „method B“), and a combination of sonication and bead-beating (designation „method C“).

**Method S** (water filter and seagrass leaves only):

- cells were dislodged by manual swabbing both sides of the leaves, the water filter respectively with a sterile cotton swab
- swab was added to a 2 mL reaction tube containing 2 mL of PBS, reaction tube was centrifuged at 10,000 x g for 10 min and the supernatant and swab were discarded
- the remaining pellet was resuspended in CD1 – solution of the DNeasy PowerSoil Kit and extraction proceeded as described in the main text

**Method B** (seagrass leaves and -roots only):

- as described in the main text, but without sonication

**Method C** (all sample types)

- as described in the main text

To test whether cell dislodgement method had an influence on community composition, the quality filtered ASV tables (16S and 18S) that resulted from amplicon sequencing were tested via PERMANOVA based on Bray-Curtis-Dissimilarities (adonis2- function of the vegan-package) and ‘method’ as explanatory variable.

**Supplementary table S1:** Overview of samples and the corresponding cell detachment procedure before DNA extraction. Method S = “Swabbing”, Method B = “Bead Beating”, Method C = “Sonication & Bead Beating”.

| sample | Method | sample | Method | sample | Method |
|---|---|---|---|---|---|
| G-H8-RO-2 | S | G-H2-RO-2 | B | G-H1-WF | C |
| G-H8-SGO-3 | S | G-H2-RO-3 | B | G-H2-RO-1 | C |
| G-H8-SGY-2 | S | G-H2-SGO-2 | B | G-H2-RO-4 | C |
| G-H8-WF | S | G-H2-SGO-3 | B | G-H2-SGO-1 | C |
| | | G-H2-SGY-2 | B | G-H2-SGO-4 | C |
| | | G-H2-SGY-3 | B | G-H2-SGY-1 | C |
| | | G-H9-RO-2 | B | G-H2-SGY-4 | C |
| | | G-H9-RO-3 | B | G-H2-SS | C |
| | | G-H9-SGO-2 | B | G-H2-WF | C |
| | | G-H9-SGO-3 | B | G-H3-WF | C |
| | | G-H9-SGY-2 | B | G-H6-WF | C |
| | | G-H9-SGY-3 | B | G-H7-SS | C |
| | | | | G-H7-WF | C |
| | | | | G-H8-RO-1 | C |
| | | | | G-H8-RO-3 | C |
| | | | | G-H8-RO-4 | C |
| | | | | G-H8-SGO-1 | C |
| | | | | G-H8-SGO-2 | C |
| | | | | G-H8-SGO-4 | C |
| | | | | G-H8-SGY-1 | C |
| | | | | G-H8-SGY-3 | C |
| | | | | G-H8-SGY-4 | C |
| | | | | G-H8-SS | C |
| | | | | G-H9-RO-1 | C |
| | | | | G-H9-RO-4 | C |
| | | | | G-H9-SGO-1 | C |
| | | | | G-H9-SGO-4 | C |
| | | | | G-H9-SGY-1 | C |
| | | | | G-H9-SGY-4 | C |
| | | | | G-H9-SS | C |
| | | | | G-H9-WF | C |

**Supplementary table S2**: Overview of picked colonies grown on selective Agar per sample type with number of colonies identified as *Vibrio* spp. and other bacteria (non-*Vibrio*) based on sequencing of the hypervariable V3-V4 regions of the 16S rRNA gene and correction factor as the ratio of non-*Vibrio* to *Vibrio* spp.

| Sampling day | Sample type | Picked colonies | Colonies identified as *Vibrio* spp. | Non *Vibrio* colonies | % *Vibrio* spp. | corr.faktor (ratio non-*Vibrio*/ *Vibrio*) |
|---|---|---|---|---|---|---|
| 1 | water | 19 | 17 | 2 | 89.47 | 0.11 |
| | seagrass | 23 | 23 | 0 | 100 | 0.00 |
| | sediment | 15 | 11 | 4 | 73.33 | 0.27 |
| 2 | water | 23 | 14 | 9 | 60.87 | 0.39 |
| | seagrass | 42 | 37 | 5 | 88.10 | 0.12 |
| | sediment | 11 | 8 | 3 | 72.73 | 0.27 |

**Supplementary table S3:** Differentially abundant genera on seagrass roots compared to young seagrass leaves. A positive log2 fold change indicates a higher abundance on roots than on young leaves, a negative indicates less abundance. BaseMean = mean of normalized counts for all samples, lfcSE = log2 fold change standard error, stat = Wald statistic, pvalue = Wald test p-value, padj = p value adjusted for multiple testing (Benjamini-Hochberg).

| Genus | baseMean | log2 fold change | lfcSE | stat | pvalue | padj |
|---|---|---|---|---|---|---|
| Amphritea | 735.918876 | 10.3070222 | 1.12823612 | 9.13551867 | 6.51E-20 | 3.61E-18 |
| Aquimonas | 3.9047755 | 1.39439653 | 0.6093033 | 2.28850974 | 0.02210785 | 0.03718139 |
| Arcobacter | 2279.7208 | 8.96211114 | 0.88178959 | 10.1635484 | 2.88E-24 | 6.40E-22 |
| Bdellovibrio | 10.4749279 | -2.1945898 | 0.8587975 | -2.5554218 | 0.01060592 | 0.01995351 |
| Ca. Omnitrophus | 3.77358584 | 3.29178863 | 0.90978181 | 3.61821769 | 0.00029664 | 0.0008031 |
| Calothrix | 73.9823314 | -3.7708144 | 0.61919818 | -6.0898344 | 1.13E-09 | 6.78E-09 |
| Candidatus Riegeria | 34.179361 | 8.18960688 | 1.15513838 | 7.0897193 | 1.34E-12 | 1.24E-11 |
| Carboxylicivirga | 46.191213 | 8.01228047 | 1.13969402 | 7.03020312 | 2.06E-12 | 1.76E-11 |
| Cellvibrio | 19.6309667 | 7.74209174 | 1.05091075 | 7.36703068 | 1.74E-13 | 2.04E-12 |
| Desulfobacter | 11.2309764 | 5.25100982 | 0.97105768 | 5.40751586 | 6.39E-08 | 2.90E-07 |
| Desulfobulbus | 135.560385 | 7.2957478 | 0.90233083 | 8.08544669 | 6.19E-16 | 1.25E-14 |
| Desulfoconvexum | 26.1328141 | 6.56336002 | 1.03481307 | 6.3425562 | 2.26E-10 | 1.57E-09 |
| Desulfomonile | 8.24617772 | 4.56023932 | 0.90157407 | 5.05808614 | 4.23E-07 | 1.71E-06 |
| Desulfopila | 32.94306 | 8.69059723 | 1.06784582 | 8.13843825 | 4.00E-16 | 8.89E-15 |
| Desulfovibrio | 172.03018 | 6.28862524 | 1.0670362 | 5.89354442 | 3.78E-09 | 2.00E-08 |
| Erythrobacter | 140.656138 | -1.8812889 | 0.66276775 | -2.8385342 | 0.00453213 | 0.00931604 |
| Flexibacter | 6.3774194 | 3.99136811 | 1.12497473 | 3.54796245 | 0.00038822 | 0.00101395 |
| Fluviicola | 183.967057 | 1.01993473 | 0.72573988 | 1.4053723 | 0.1599106 | 0.22049784 |
| Fusibacter | 171.123699 | 7.75651621 | 0.92188675 | 8.41374084 | 3.97E-17 | 1.41E-15 |
| Glaciecola | 615.17371 | -3.7795138 | 1.23316854 | -3.0648801 | 0.00217757 | 0.00483421 |
| Haliangium | 4.20306774 | 1.31316838 | 0.78942367 | 1.663452 | 0.09622199 | 0.14240855 |
| Hirschia | 242.526682 | -1.2162524 | 1.13320882 | -1.0732818 | 0.28314469 | 0.35116269 |
| Jatrophihabitans | 6.43659299 | 2.75860577 | 0.85949053 | 3.20958251 | 0.00132928 | 0.00320761 |
| Labrenzia | 41.9294005 | 7.70226692 | 0.97847922 | 7.87167145 | 3.50E-15 | 5.98E-14 |
| Lachnoclostridium 5 | 4.35423051 | 4.11537233 | 0.98804689 | 4.16515894 | 3.11E-05 | 0.00010465 |
| Lachnotalea | 6.01217531 | 5.5781197 | 1.05708398 | 5.2768936 | 1.31E-07 | 5.82E-07 |
| Lewinella | 121.550085 | -0.9033634 | 0.68641147 | -1.3160669 | 0.18815159 | 0.24862889 |
| Litorimicrobium | 1122.07546 | 0.44898037 | 1.09994308 | 0.40818509 | 0.68313779 | 0.72911822 |
| Lutibacter | 48.08356 | 6.18757566 | 1.05118788 | 5.88627 | 3.95E-09 | 2.04E-08 |
| Lutimonas | 14.7129471 | 4.33682096 | 0.77779679 | 5.57577636 | 2.46E-08 | 1.14E-07 |
| Methanolobus | 7.40689709 | 5.48966944 | 1.05680386 | 5.19459634 | 2.05E-07 | 8.59E-07 |
| Methylotenera | 3252.2544 | 1.52310971 | 0.79744268 | 1.90999271 | 0.05613415 | 0.08901273 |
| MWH-UniP1 aquatic group | 1.77098221 | -0.5223825 | 0.70814474 | -0.7376776 | 0.46071036 | 0.52993627 |
| Neptunomonas | 16.5591997 | 7.008529 | 1.09394808 | 6.40663767 | 1.49E-10 | 1.07E-09 |
| Oceaniovalibus | 72.2900302 | -0.6749533 | 0.83343949 | -0.8098407 | 0.41803171 | 0.49363319 |
| Oleiphilus | 6.98088652 | 4.41633835 | 0.92150086 | 4.79254938 | 1.65E-06 | 6.09E-06 |

| | | | | | | |
|---|---|---|---|---|---|---|
| Opitutus | 13.278746 | 5.0720749 | 0.87555299 | 5.79299595 | 6.91E-09 | 3.49E-08 |
| Pacificibacter | 1123.77174 | -1.2086202 | 1.01268538 | -1.1934805 | 0.23268125 | 0.3056523 |
| Paludibacter | 48.1768817 | 7.67137983 | 1.29937933 | 5.90388016 | 3.55E-09 | 1.92E-08 |
| Pelagicoccus | 236.494092 | 6.19609921 | 0.73756278 | 8.40077534 | 4.44E-17 | 1.41E-15 |
| Pelobacter | 61.4665332 | 6.91395597 | 1.22312874 | 5.6526805 | 1.58E-08 | 7.62E-08 |
| Perspicuibacter | 36.6038777 | -2.2284169 | 1.02085475 | -2.1828932 | 0.02904367 | 0.04847891 |
| Phaselicystis | 11.4912443 | 4.51946089 | 0.98441902 | 4.59099307 | 4.41E-06 | 1.58E-05 |
| Propionigenium | 4.08425504 | 3.09734634 | 0.79180591 | 3.91174945 | 9.16E-05 | 0.00026079 |
| Propionivibrio | 91.8485626 | 8.60988871 | 1.18219922 | 7.28294229 | 3.27E-13 | 3.63E-12 |
| Pseudahrensia | 23.1978165 | -0.5753297 | 0.62257349 | -0.9241154 | 0.35542624 | 0.42651149 |
| Psychromonas | 13.6305086 | 6.10112672 | 0.97961505 | 6.22808595 | 4.72E-10 | 2.99E-09 |
| Puniceicoccus | 143.849478 | 8.37674859 | 1.01153673 | 8.28121054 | 1.22E-16 | 3.38E-15 |
| Rubidimonas | 105.267461 | -2.9977272 | 1.03165698 | -2.9057402 | 0.00366386 | 0.00767336 |
| Saccharicrinis | 188.995862 | 7.28884457 | 1.19053024 | 6.12235147 | 9.22E-10 | 5.69E-09 |
| Sedimenticola | 68.3017118 | 8.25632877 | 1.08687215 | 7.5964121 | 3.04E-14 | 4.22E-13 |
| Shewanella | 197.993297 | 6.77373374 | 0.90691785 | 7.46896064 | 8.08E-14 | 9.97E-13 |
| Simiduia | 91.1609297 | 5.42839541 | 0.86808122 | 6.25332665 | 4.02E-10 | 2.62E-09 |
| Spirochaeta | 158.645381 | 7.57949007 | 0.85969148 | 8.8165234 | 1.18E-18 | 5.24E-17 |
| Sulfurimonas | 4190.6909 | 8.21159798 | 1.08767374 | 7.54968854 | 4.36E-14 | 5.70E-13 |
| Sulfurospirillum | 258.418019 | 7.71897172 | 1.11547755 | 6.91988084 | 4.52E-12 | 3.58E-11 |
| Teredinibacter | 57.3542972 | 6.23912113 | 0.88517877 | 7.0484306 | 1.81E-12 | 1.61E-11 |
| Thalassospira | 13.913719 | 6.79007771 | 1.01544442 | 6.68680388 | 2.28E-11 | 1.69E-10 |
| Thioalkalispira | 50.6171978 | 6.88047074 | 1.08917984 | 6.31711171 | 2.66E-10 | 1.79E-09 |
| Thiohalocapsa | 2.10510796 | 2.55369845 | 0.80610917 | 3.16793127 | 0.00153528 | 0.00358258 |
| Uliginosibacterium | 350.077909 | 8.69552103 | 0.8896915 | 9.77363617 | 1.46E-22 | 1.62E-20 |
| Unclassified | 28735.8281 | 1.86743806 | 0.25929209 | 7.20206346 | 5.93E-13 | 6.27E-12 |
| Unknown Caldithrix family incertae sedis genus 1 | 5.57743773 | 5.14894572 | 0.98693544 | 5.21710488 | 1.82E-07 | 7.76E-07 |
| Unknown Defluviitaleaceae genus 1 | 118.999741 | 9.09815716 | 1.11770479 | 8.14003593 | 3.95E-16 | 8.89E-15 |
| Unknown Defluviitaleaceae genus 2 | 205.756591 | 9.94501964 | 1.2994904 | 7.65301508 | 1.96E-14 | 2.91E-13 |
| Unknown Defluviitaleaceae genus 3 | 110.449849 | 9.73713263 | 1.21199856 | 8.03394734 | 9.44E-16 | 1.75E-14 |
| Unknown Desulfobacteraceae genus 2 | 30.5732102 | 7.7806993 | 1.12697321 | 6.90406765 | 5.05E-12 | 3.87E-11 |
| Unknown Desulfovibrionaceae genus 2 | 15.4388743 | 5.55078615 | 1.0525487 | 5.27366207 | 1.34E-07 | 5.82E-07 |
| Unknown Lweinellacea genus 2 | 17.807388 | 2.88690365 | 1.04261092 | 2.76891753 | 0.00562429 | 0.01145497 |

| | | | | | | |
|---|---|---|---|---|---|---|
| Unknown Lweinellacea genus 3 | 3.18328626 | 1.07257503 | 0.78217368 | 1.37127477 | 0.17028932 | 0.23192778 |
| Unknown Planctomycetaceae genus 4 | 2.5585551 | -2.8424593 | 0.89800585 | -3.1653016 | 0.00154922 | 0.00358258 |
| Unknown Planctomycetaceae genus 8 | 1.65896788 | 1.4388219 | 0.69143472 | 2.08092227 | 0.03744102 | 0.06156968 |
| Unknown Prolixibacteraceae genus 2 | 76.2231348 | 7.94908404 | 1.14076777 | 6.96818777 | 3.21E-12 | 2.64E-11 |
| Unknown Saprospiraceae genus 1 | 184.30297 | -6.571846 | 1.59720276 | -4.1145972 | 3.88E-05 | 0.00012662 |
| Vibrio | 557.280308 | 8.5155976 | 0.93163678 | 9.1404695 | 6.22E-20 | 3.61E-18 |

**Supplementary table S4:** Differentially abundant genera on seagrass roots compared to old seagrass leaves. A positive log2 fold change indicates a higher abundance on roots than on old leaves, a negative indicates less abundance. BaseMean = mean of normalized counts for all samples, lfcSE = log2 fold change standard error, stat = Wald statistic, pvalue = Wald test p-value, padj = p value adjusted for multiple testing (Benjamini-Hochberg).

| Genus | baseMean | log2 fold change | lfcSE | stat | pvalue | padj |
|---|---|---|---|---|---|---|
| Acholeplasma | 5.93673098 | 4.090894631 | 1.10214704 | 3.71175033 | 0.00020583 | 0.00057118 |
| Actibacter | 24.1283673 | 3.20661021 | 0.81940072 | 3.91336023 | 9.10E-05 | 0.00026079 |
| Aestuariibacter | 5.79326439 | 3.562339542 | 0.87883177 | 4.05349426 | 5.05E-05 | 0.00015558 |
| Amphritea | 735.918876 | 10.30702217 | 1.12823612 | 9.13551867 | 6.51E-20 | 3.61E-18 |
| Aquimonas | 3.9047755 | 1.394396526 | 0.6093033 | 2.28850974 | 0.02210785 | 0.03718139 |
| Arcobacter | 2279.7208 | 8.962111144 | 0.88178959 | 10.1635484 | 2.88E-24 | 6.40E-22 |
| Bdellovibrio | 10.4749279 | -2.19458982 | 0.8587975 | -2.5554218 | 0.01060592 | 0.01995351 |
| Blastopirellula | 214.706373 | -1.982251475 | 0.55552743 | -3.5682333 | 0.0003594 | 0.00094983 |
| Ca. Omnitrophus | 3.77358584 | 3.291788626 | 0.90978181 | 3.61821769 | 0.00029664 | 0.0008031 |
| Caldithrix | 31.6502803 | 4.265156831 | 0.88571428 | 4.81549968 | 1.47E-06 | 5.52E-06 |
| Calothrix | 73.9823314 | -3.770814369 | 0.61919818 | -6.0898344 | 1.13E-09 | 6.78E-09 |
| Candidatus Accumulibacter | 3.15277187 | 3.540150645 | 0.85200005 | 4.15510616 | 3.25E-05 | 0.00010773 |
| Candidatus Amoebophilus | 32.191261 | -1.89089455 | 0.70827311 | -2.6697252 | 0.00759134 | 0.0150127 |
| Candidatus Anammoximicrobium | 3.43412558 | 1.986884529 | 0.80514442 | 2.46773683 | 0.01359702 | 0.02494661 |
| Candidatus Microthrix | 34.707504 | 1.776593328 | 0.68103631 | 2.6086617 | 0.00908971 | 0.01739582 |

| | | | | | | |
|---|---|---|---|---|---|---|
| Candidatus Riegeria | 34.179361 | 8.189606877 | 1.15513838 | 7.0897193 | 1.34E-12 | 1.24E-11 |
| Carboxylicivirga | 46.191213 | 8.012280468 | 1.13969402 | 7.03020312 | 2.06E-12 | 1.76E-11 |
| Cellvibrio | 19.6309667 | 7.742091742 | 1.05091075 | 7.36703068 | 1.74E-13 | 2.04E-12 |
| Clostridium sensu stricto | 18.4269132 | 2.645118671 | 0.78537559 | 3.36796647 | 0.00075725 | 0.00191033 |
| Colwellia | 9.5291246 | 2.2043316 | 0.96014489 | 2.29583225 | 0.02168547 | 0.03718139 |
| Crocinitomix | 14.650321 | 1.225716008 | 0.53489312 | 2.29151576 | 0.0219336 | 0.03718139 |
| Defluviitaleaceae UCG-011 | 51.6577337 | 7.972520174 | 1.11097496 | 7.17614751 | 7.17E-13 | 7.24E-12 |
| Desulfatiglans | 26.1418331 | 4.03493609 | 0.8827896 | 4.57066564 | 4.86E-06 | 1.71E-05 |
| Desulfobacter | 11.2309764 | 5.25100982 | 0.97105768 | 5.40751586 | 6.39E-08 | 2.90E-07 |
| Desulfobulbus | 135.560385 | 7.295747804 | 0.90233083 | 8.08544669 | 6.19E-16 | 1.25E-14 |
| Desulfococcus | 10.2304227 | 3.929598905 | 0.96664344 | 4.06519999 | 4.80E-05 | 0.0001522 |
| Desulfoconvexum | 26.1328141 | 6.563360019 | 1.03481307 | 6.3425562 | 2.26E-10 | 1.57E-09 |
| Desulfomonile | 8.24617772 | 4.56023932 | 0.90157407 | 5.05808614 | 4.23E-07 | 1.71E-06 |
| Desulfopila | 32.94306 | 8.690597233 | 1.06784582 | 8.13843825 | 4.00E-16 | 8.89E-15 |
| Desulforhopalus | 318.585147 | 7.303227622 | 1.01857525 | 7.17004233 | 7.50E-13 | 7.24E-12 |
| Desulfosarcina | 130.368062 | 4.383859621 | 0.98718893 | 4.4407504 | 8.96E-06 | 3.11E-05 |
| Desulfovibrio | 172.03018 | 6.288625238 | 1.0670362 | 5.89354442 | 3.78E-09 | 2.00E-08 |
| Desulfuromonas | 10.387281 | 5.004109022 | 1.20122436 | 4.16584044 | 3.10E-05 | 0.00010465 |
| Devosia | 8.363513 | 2.774092991 | 0.92889654 | 2.98643916 | 0.00282247 | 0.00604653 |
| Draconibacterium | 32.6405635 | 5.625818279 | 0.94987476 | 5.92269479 | 3.17E-09 | 1.76E-08 |
| Ekhidna | 19.855581 | -2.90763628 | 0.96988358 | -2.997923 | 0.00271826 | 0.00591622 |
| Erythrobacter | 140.656138 | -1.881288915 | 0.66276775 | -2.8385342 | 0.00453213 | 0.00931604 |
| Flaviramulus | 5.94984384 | 3.770128858 | 0.92891876 | 4.05862064 | 4.94E-05 | 0.00015435 |
| Flavobacterium | 122.052621 | 3.050225559 | 0.77725894 | 3.9243364 | 8.70E-05 | 0.00025404 |
| Flexibacter | 6.3774194 | 3.991368112 | 1.12497473 | 3.54796245 | 0.00038822 | 0.00101395 |
| Fodinicola | 62.249617 | 2.095158926 | 0.78458175 | 2.67041505 | 0.00757575 | 0.0150127 |
| Fusibacter | 171.123699 | 7.756516208 | 0.92188675 | 8.41374084 | 3.97E-17 | 1.41E-15 |
| Gemmata | 3.82035573 | -2.448348485 | 0.95056358 | -2.5756809 | 0.01000429 | 0.0189825 |
| Gemmatimonas | 5.93946942 | -1.729747235 | 0.72719295 | -2.3786634 | 0.01737554 | 0.03069157 |
| Glaciecola | 615.17371 | -3.779513766 | 1.23316854 | -3.0648801 | 0.00217757 | 0.00483421 |
| Granulosicoccus | 480.848188 | -2.237423509 | 0.87717996 | -2.5507007 | 0.01075066 | 0.02005586 |
| Ignavibacterium | 8.47369664 | 2.944937842 | 0.96047405 | 3.06612952 | 0.00216849 | 0.00483421 |
| Jatrophihabitans | 6.43659299 | 2.758605771 | 0.85949053 | 3.20958251 | 0.00132928 | 0.00320761 |
| Labrenzia | 41.9294005 | 7.702266922 | 0.97847922 | 7.87167145 | 3.50E-15 | 5.98E-14 |
| Lachnoclostridium 5 | 4.35423051 | 4.115372327 | 0.98804689 | 4.16515894 | 3.11E-05 | 0.00010465 |
| Lachnotalea | 6.01217531 | 5.578119699 | 1.05708398 | 5.2768936 | 1.31E-07 | 5.82E-07 |
| Leptolyngbya | 36.1026784 | -2.801499157 | 0.68601233 | -4.0837446 | 4.43E-05 | 0.00014258 |
| Limnobacter | 48.0472133 | -1.737192154 | 0.49972379 | -3.4763047 | 0.00050837 | 0.00131232 |
| Lutibacter | 48.08356 | 6.187575663 | 1.05118788 | 5.88627 | 3.95E-09 | 2.04E-08 |
| Lutimonas | 14.7129471 | 4.336820962 | 0.77779679 | 5.57577636 | 2.46E-08 | 1.14E-07 |
| Maribacter | 38.6276433 | -2.55875493 | 0.92854212 | -2.7556692 | 0.00585722 | 0.01182094 |
| Marinicella | 6.30633097 | 3.50387775 | 0.89113779 | 3.93191466 | 8.43E-05 | 0.00024945 |
| Marinomonas | 383.812958 | 4.109392436 | 0.79526103 | 5.16735041 | 2.37E-07 | 9.76E-07 |
| Methanolobus | 7.40689709 | 5.489669443 | 1.05680386 | 5.19459634 | 2.05E-07 | 8.59E-07 |
| Mobilitalea | 5.89487758 | 5.122390693 | 1.04048625 | 4.92307387 | 8.52E-07 | 3.32E-06 |

| | | | | | | |
|---|---|---|---|---|---|---|
| Neptunomonas | 16.5591997 | 7.008528996 | 1.09394808 | 6.40663767 | 1.49E-10 | 1.07E-09 |
| Novosphingobium | 3.82584348 | 1.832321357 | 0.77716914 | 2.3576867 | 0.01838921 | 0.03214491 |
| Oleiphilus | 6.98088652 | 4.416338352 | 0.92150086 | 4.79254938 | 1.65E-06 | 6.09E-06 |
| OM60(NOR5) clade | 2.28105258 | 1.994255145 | 0.83143191 | 2.39857902 | 0.01645882 | 0.02970617 |
| Opitutus | 13.278746 | 5.072074905 | 0.87555299 | 5.79299595 | 6.91E-09 | 3.49E-08 |
| Paludibacter | 48.1768817 | 7.671379828 | 1.29937933 | 5.90388016 | 3.55E-09 | 1.92E-08 |
| Paraglaciecola | 787.720831 | 3.827074576 | 0.81188107 | 4.71383642 | 2.43E-06 | 8.85E-06 |
| Parvularcula | 7.6728115 | -2.969259944 | 0.92787871 | -3.2000518 | 0.00137403 | 0.00327994 |
| Pelagicoccus | 236.494092 | 6.196099208 | 0.73756278 | 8.40077534 | 4.44E-17 | 1.41E-15 |
| Pelobacter | 61.4665332 | 6.913955965 | 1.22312874 | 5.6526805 | 1.58E-08 | 7.62E-08 |
| Perspicuibacter | 36.6038777 | -2.228416946 | 1.02085475 | -2.1828932 | 0.02904367 | 0.04847891 |
| Phaselicystis | 11.4912443 | 4.519460894 | 0.98441902 | 4.59099307 | 4.41E-06 | 1.58E-05 |
| Pirellula | 187.575708 | -2.058036266 | 0.61775441 | -3.3314797 | 0.00086386 | 0.00215479 |
| Planctomyces | 140.783043 | -1.396212502 | 0.43784416 | -3.1888343 | 0.00142848 | 0.00337364 |
| Propionigenium | 4.08425504 | 3.097346343 | 0.79180591 | 3.91174945 | 9.16E-05 | 0.00026079 |
| Propionivibrio | 91.8485626 | 8.609888714 | 1.18219922 | 7.28294229 | 3.27E-13 | 3.63E-12 |
| Pseudoalteromonas | 217.200437 | 5.069007161 | 1.04484582 | 4.85144034 | 1.23E-06 | 4.69E-06 |
| Pseudohaliea | 6.12413328 | -3.802136927 | 1.05252048 | -3.6124114 | 0.00030336 | 0.0008114 |
| Pseudomonas | 56.2796399 | 5.661293459 | 1.01318826 | 5.58760271 | 2.30E-08 | 1.09E-07 |
| Psychromonas | 13.6305086 | 6.101126724 | 0.97961505 | 6.22808595 | 4.72E-10 | 2.99E-09 |
| Puniceicoccus | 143.849478 | 8.376748587 | 1.01153673 | 8.28121054 | 1.22E-16 | 3.38E-15 |
| Reinekea | 113.336401 | 7.128389836 | 0.9302942 | 7.66251135 | 1.82E-14 | 2.89E-13 |
| Rheinheimera | 137.159211 | 4.56650184 | 0.76254427 | 5.98850722 | 2.12E-09 | 1.21E-08 |
| Rhizobium | 6.6950033 | 2.732854548 | 0.90779018 | 3.01044737 | 0.00260863 | 0.00573382 |
| Rhodopirellula | 132.211979 | -1.70793884 | 0.74581597 | -2.2900272 | 0.02201974 | 0.03718139 |
| Roseivirga | 8.36014301 | -2.682021217 | 1.01851922 | -2.6332554 | 0.00845707 | 0.01632583 |
| Rubidimonas | 105.267461 | -2.997727161 | 1.03165698 | -2.9057402 | 0.00366386 | 0.00767336 |
| Rubrivirga | 12.266633 | -2.310855566 | 0.91228512 | -2.5330409 | 0.01130777 | 0.02091938 |
| Ruminococcus 1 | 2.68846561 | 3.207545885 | 0.85997837 | 3.72979832 | 0.00019163 | 0.00053851 |
| Saccharicrinis | 188.995862 | 7.288844575 | 1.19053024 | 6.12235147 | 9.22E-10 | 5.69E-09 |
| Sedimenticola | 68.3017118 | 8.256328766 | 1.08687215 | 7.5964121 | 3.04E-14 | 4.22E-13 |
| SEEP-SRB1 | 23.4330409 | 3.172430823 | 0.97239408 | 3.26249501 | 0.00110436 | 0.00272409 |
| Shewanella | 197.993297 | 6.773733743 | 0.90691785 | 7.46896064 | 8.08E-14 | 9.97E-13 |
| Simiduia | 91.1609297 | 5.428395408 | 0.86808122 | 6.25332665 | 4.02E-10 | 2.62E-09 |
| SM1A02 | 23.7157193 | -2.017656763 | 0.84856403 | -2.3777307 | 0.01741954 | 0.03069157 |
| Spirochaeta | 158.645381 | 7.57949007 | 0.85969148 | 8.8165234 | 1.18E-18 | 5.24E-17 |
| Sporichthya | 3.85500091 | 1.869176421 | 0.79799409 | 2.34234368 | 0.01916306 | 0.03323593 |
| Sulfurimonas | 4190.6909 | 8.211597984 | 1.08767374 | 7.54968854 | 4.36E-14 | 5.70E-13 |
| Sulfurospirillum | 258.418019 | 7.718971723 | 1.11547755 | 6.91988084 | 4.52E-12 | 3.58E-11 |
| Sulfurovum | 77.205784 | 4.24414898 | 1.16217895 | 3.65188939 | 0.00026032 | 0.00071346 |
| Sva0081 sediment group | 142.040798 | 2.566457646 | 0.96211798 | 2.66750825 | 0.0076416 | 0.0150127 |
| Teredinibacter | 57.3542972 | 6.239121129 | 0.88517877 | 7.0484306 | 1.81E-12 | 1.61E-11 |
| Thalassospira | 13.913719 | 6.790077706 | 1.01544442 | 6.68680388 | 2.28E-11 | 1.69E-10 |
| Thalassotalea | 84.6253277 | 3.823964598 | 1.60373 | 2.3844192 | 0.0171061 | 0.03062544 |
| Thioalkalispira | 50.6171978 | 6.880470738 | 1.08917984 | 6.31711171 | 2.66E-10 | 1.79E-09 |
| Thiohalocapsa | 2.10510796 | 2.553698448 | 0.80610917 | 3.16793127 | 0.00153528 | 0.00358258 |

| | | | | | | |
|---|---|---|---|---|---|---|
| Thiothrix | 82.4664598 | 2.912724909 | 0.97567531 | 2.98534243 | 0.00283261 | 0.00604653 |
| Truepera | 29.7318538 | -1.959731663 | 0.49740288 | -3.9399283 | 8.15E-05 | 0.00024452 |
| Uliginosibacterium | 350.077909 | 8.695521035 | 0.8896915 | 9.77363617 | 1.46E-22 | 1.62E-20 |
| Unclassified | 28735.8281 | 1.867438061 | 0.25929209 | 7.20206346 | 5.93E-13 | 6.27E-12 |
| Unknown Caldithrix family incertae sedis genus 1 | 5.57743773 | 5.148945719 | 0.98693544 | 5.21710488 | 1.82E-07 | 7.76E-07 |
| Unknown Caldithrix family incertae sedis genus 2 | 8.73805286 | 3.967881635 | 0.99442235 | 3.99013724 | 6.60E-05 | 0.00020082 |
| Unknown Defluviit aleaceae genus 1 | 118.999741 | 9.09815716 | 1.11770479 | 8.14003593 | 3.95E-16 | 8.89E-15 |
| Unknown Defluviit aleaceae genus 2 | 205.756591 | 9.945019641 | 1.2994904 | 7.65301508 | 1.96E-14 | 2.91E-13 |
| Unknown Defluviit aleaceae genus 3 | 110.449849 | 9.737132626 | 1.21199856 | 8.03394734 | 9.44E-16 | 1.75E-14 |
| Unknown Defluviit aleaceae genus 4 | 46.3573459 | 7.45944527 | 1.48542067 | 5.02177289 | 5.12E-07 | 2.03E-06 |
| Unknown Desulfo bacteraceae genu s 1 | 32.1365956 | 7.327036933 | 1.20464401 | 6.08232546 | 1.18E-09 | 6.92E-09 |
| Unknown Desulfo bacteraceae genu s 2 | 30.5732102 | 7.780699296 | 1.12697321 | 6.90406765 | 5.05E-12 | 3.87E-11 |
| Unknown Desulfov ibrionaceae genus 2 | 15.4388743 | 5.550786147 | 1.0525487 | 5.27366207 | 1.34E-07 | 5.82E-07 |
| Unknown Lweinell acea genus 2 | 17.807388 | 2.886903652 | 1.04261092 | 2.76891753 | 0.00562429 | 0.01145497 |
| Unknown Plancto mycetaceae genu s 1 | 8.47666495 | -2.050605693 | 0.62958968 | -3.257051 | 0.00112576 | 0.00274636 |
| Unknown Plancto mycetaceae genu s 2 | 4.02840541 | 2.03210482 | 0.71540424 | 2.84049871 | 0.00450431 | 0.00931604 |
| Unknown Plancto mycetaceae genu s 20 | 2.71054464 | -2.572187696 | 0.97596187 | -2.6355412 | 0.00840032 | 0.01632583 |
| Unknown Plancto mycetaceae genu s 3 | 6.09126861 | 2.486154217 | 0.80845138 | 3.07520559 | 0.00210357 | 0.00476524 |

| | | | | | | |
|---|---|---|---|---|---|---|
| Unknown Planctomycetaceae genus 4 | 2.5585551 | -2.842459332 | 0.89800585 | -3.1653016 | 0.00154922 | 0.00358258 |
| Unknown Prolixibacteraceae genus 1 | 42.5638975 | 6.59366678 | 1.15964323 | 5.68594425 | 1.30E-08 | 6.42E-08 |
| Unknown Prolixibacteraceae genus 2 | 76.2231348 | 7.949084042 | 1.14076777 | 6.96818777 | 3.21E-12 | 2.64E-11 |
| Unknown Rhodospirillales Incertae Sedis genus 1 | 3.21711461 | -2.652894201 | 0.90115756 | -2.9438739 | 0.00324132 | 0.00685308 |
| Unknown Saprospiraceae genus 1 | 184.30297 | -6.571845972 | 1.59720276 | -4.1145972 | 3.88E-05 | 0.00012662 |
| Unknown Sphingomonadaceae genus 1 | 3.59842373 | -2.884688133 | 0.8502376 | -3.3928024 | 0.00069182 | 0.00176532 |
| Unknown Syntrophaceae genus 1 | 4.31098146 | 2.385424602 | 0.75496456 | 3.1596511 | 0.00157958 | 0.00361513 |
| Vibrio | 557.280308 | 8.515597601 | 0.93163678 | 9.1404695 | 6.22E-20 | 3.61E-18 |
| Winogradskyella | 7.92145517 | 1.866786202 | 0.76490566 | 2.44054437 | 0.01466514 | 0.02668575 |
